# HNRNPK is retained in the cytoplasm by Keratin 19 to stabilize target mRNAs

**DOI:** 10.1101/2022.01.24.477557

**Authors:** Arwa Fallatah, Dimitrios G. Anastasakis, Amirhossein Manzourolajdad, Pooja Sharma, Xiantao Wang, Alexis Jacob, Sarah Alsharif, Ahmed Elgerbi, Pierre A. Coulombe, Markus Hafner, Byung Min Chung

## Abstract

Heterogeneous nuclear ribonucleoprotein K (HNRNPK) regulates pre-mRNA processing and long non-coding RNA localization in the nucleus. It was previously shown that shuttling of HNRNPK to the cytoplasm promotes cell proliferation and cancer metastasis. However, the mechanism of HNRNPK cytoplasmic localization, its cytoplasmic RNA ligands, and impact on posttranscriptional gene regulation remain uncharacterized. Here we show that the intermediate filament protein Keratin 19 (K19) directly interacts with HNRNPK and sequesters it in the cytoplasm. Correspondingly, in K19 knockout breast cancer cells, HNRNPK does not localize in the cytoplasm, resulting in reduced cell proliferation. We mapped cytoplasmic HNRNPK target mRNAs using PAR-CLIP where transcriptome data to show that, in the cytoplasm, HNRNPK stabilizes target mRNAs bound to the 3’ untranslated region at the expected C-rich sequence elements. Furthermore, these mRNAs are typically involved in cancer progression and include the p53 signaling pathway that is dysregulated upon HNRNPK knockdown or K19 knockout. This study identifies how a cytoskeletal protein can directly regulate gene expression by controlling subcellular localization of RNA binding proteins to support pathways involved in cancer progression.

## Introduction

Heterogeneous nuclear ribonucleoproteins (hnRNP) are a group of abundant nuclear RNA binding proteins (RBPs) consisting of more than 20 members from various RBP families that predominantly localize to the cell nucleus(1). It comprises three K-homology (KH) RNA binding domains with clear preference to cytosine-rich sequences(2), in addition to a less characterized K interactive region (KI) that is thought to allow interaction with multiple other proteins. HNRNPK plays crucial roles in several biological processes, including development, axonal outgrowth, cell proliferation, and migration(3–6). In cancer, overexpression of HNRNPK promotes tumor progression and correlates with poor patient survival(7), likely by directly affecting the expression and activities of oncogenes and tumor suppressors such as EIF4E(8), c-MYC(9), c-SRC(10) and MDM2(11). As the list of oncogenes and tumor suppressors regulated by HNRNPK is still growing, a comprehensive perspective of HNRNPK-target genes and its regulatory mechanism remains elusive. Considering that KH domains typically only recognize few nucleotides in RNA(12), systems-wide methods are required for the identification of HNRNPK-controlled posttranscriptional regulatory networks. While a few studies used crosslinking and immunoprecipitation (CLIP)-type approaches to catalogue HNRNPK target mRNAs(13), its genome-wide regulatory impact has not been fully determined yet.

While HNRNPK primarily resides in the nucleus(1) where it impacts all steps of gene expression, a cytoplasmic pool of HNRNPK has been linked to metastasis and poor prognosis in cancer(14–16). Although HNRNPK contains a nuclear shuttling domain(17), and posttranslational modifications, such as phosphorylation by ERK(18) and methylation by PRMT1(19) contribute to subcellular localization of HNRNPK, the precise mechanism by which HNRNPK is retained in the cytoplasm is still unclear. A previous study in skin cancer cells showed that a cytoskeletal protein keratin 17 (K17) is required for the cytoplasmic localization of HNRNPK(6), introducing keratin as a regulator of HNRNPK localization.

The keratin family is composed of 54 keratin genes, and expression of each keratin is tightly regulated in a tissue-, context-, and differentiation-specific manner(20). Among keratins, keratin 19 (K19) is the smallest with a very short tail domain. It is expressed in various simple and complex epithelial tissues and becomes upregulated in several cancers where it is used as a diagnostic and prognostic marker(21). Altered expression of K19 affects growth of cancer cells *in vitro* and tumors in mice(21), demonstrating its active role in cancer.

Inside the cell, keratins can interact with multiple proteins and regulate their localization. K19 has been shown to interact with various signaling molecules including β-catenin/RAC1(22), Egr1(23), HER2(24), and GSK3β(25) to regulate their subcellular localization. In this study, we explored the possibility of an interaction between K19 and HNRNPK and the indirect impact of K19 on posttranscriptional gene regulation of HNRNPK target mRNAs. We found that the direct interaction between K19 and HNRNPK was required for cytoplasmic HNRNPK localization in breast cancer cells. By using photoactivatable ribonucleoside-enhanced crosslinking and immunoprecipitation (PAR-CLIP) to identify mRNAs bound to HNRNPK along with RNA-seq data, we identified genes whose stability are affected by K19. Top pathways of genes whose stabilization depend on K19 and cytoplasmic HNRNPK were related to cancer signaling and included the p53 tumor suppressor pathway. These findings reveal that K19 regulates HNRNPK functions in gene expression through its physical interaction, which ultimately affects expression of genes involved in the p53 signaling pathway.

## Materials and Methods

### Cell culture

The MDA-MB-231 cell line was a generous gift from Dr. Zaver Bhujwalla (Johns Hopkins School of Medicine, Baltimore, MD). MDA-MB-231 cells were authenticated by short-tandem repeat profiling, service performed by ATCC. MDA-MB-231 *KRT19* KO cells generated using the CRISPR/Cas9 system and cells stably expressing GFP or GFP-*KRT19* generated using the lentiviral system were described previously(26). For MDA-MB-231 cells stably overexpressing vector or *KRT19*, lentiviral supernatants were generated using the pLenti CMV/TO V5 plasmids as described previously(26). Stable transductants were selected using hygromycin (100 μg/ml). Cells were grown in Dulbecco’s modified essential medium (DMEM; Gibco, Grand Island, NY) supplemented with 10% Fetal Bovine Serum (GE Healthcare, Logan, UT) and 1% penicillin /streptomycin (Gibco) in a humidified incubator at 5% CO_2_ and 37°C. To measure cell proliferation, 50,000 cells were plated on each well of six-well plates. Cells were counted using a hemacytometer 24, 48, 72, and 96 h after plating.

### Plasmids and siRNAs

To overexpress HNRNPK, cDNA was cloned out of pCMV6-AC *HNRNPK* (OriGene, Rockville, MD) using the primers described previously(6) and cloned into pmCherry-C1 (Takara Bio Inc., San Jose, CA). To silence HNRNPK expression, Accell Human SMARTpool *HNRNPK* siRNAs (Dharmacon, Lafayette, CO) was used. The siRNA sequences targeting *HNRNPK* are described in Table 1. AllStars negative control SI03650318 was purchased from Qiagen (Germantown, MD).

**Table 1.**
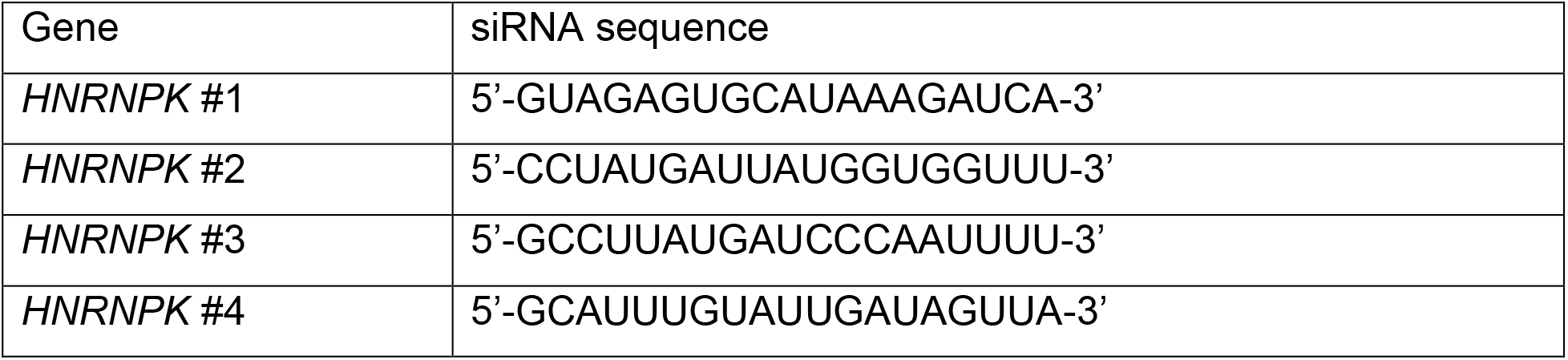
Sequences for Accell Human SMARTpool *HNRNPK* siRNAs from Dharmacon

### Transient transfection

For siRNA, RNAimax lipofectamine (Invitrogen, Waltham, MA) was used to transfect *HNRNPK* smartpool siRNA (Dharmacon) or non-targeting siRNA (Qiagen) according to the manufacturer’s protocol. For overexpression plasmids, jetOptimus DNA transfection reagent (Polyplus, Illkirch, France) was used according to the manufacturer’s protocol.

### PAR-CLIP analysis

For HNRNPK PAR-CLIP, both parental and *KRT19* KO cells were treated with 100 μM 4SU for 16 h, washed with phosphate-buffered saline and irradiated with 0.15 mJ cm^2^, 365 nm ultraviolet light in a Spectrolinker XL-1500 UV crosslinker to crosslink RNA to HNRNPK, and harvested and lysed in the equivalent of three cell pellet volumes of NP-40 lysis buffer. After spinning the lysates for 15 minutes, 10 U/ul was added to the lysates followed by the immunoprecipitation using Protein G magnetic beads. The beads were resuspended in one bead volume of dephosphorylation buffer after washing them with lysis buffer. [γ-^32^P]-ATP radiolabeling was following the dephosphorylation step. Then, the protein-RNA complexes were run on SDS-PAGE followed by autoradiography. The protein-RNA band were recovered at the corresponding size ∼55 kDa then eluted. Proteinase K digestion was performed followed by RNA recovery by acid phenol/chloroform extraction and ethanol precipitation. After RNA isolation cDNA library preparation was carried out(27). Library preparation was done using cDNA and sequences by Illumina platform. Obtained reads were processed and referenced to hg19 genome. Data analysis was performed by using PARalyzer settings where T-to-C mutation sequences were filtered.

### RNA analysis

For RNA-seq analysis, total RNAs from three biological replicates each of parental and *KRT19* KO cells. Ribosomal RNA was depleted of using the NEBNext® rRNA Depletion Kit and cDNA libraries were prepared using the NEBNext® Ultra™ Directional RNA Library Prep Kit for Illumina® (NEB, Ipswich MA). RNA was barcoded using the NEBNext Multiplex Oligos for Illumina (NEB). All samples were multiplexed and sequenced on the Illumina HiSeq 3000 platform using 50 cycles single-end sequencing. Reads were aligned to human genome version hg19 using TopHat2(28). Cufflinks and Cuffdiff were used to quantify transcripts and determine differential expression.

### Antibodies and other reagents

The following antibodies: anti-GAPDH (FL-335), anti-K19 (A-3), anti-K8 (C51), anti-K18 (C-04), anti-PARP (F-2), anti-HNRNPK (3C2) and anti-MDM2 (SMP14) were from Santa Cruz Biotechnology (Santa Cruz, CA); anti-p53 (DO-2) was from Sigma Aldrich (St. Louis, MO); and anti-GFP (12A6) was from the Developmental Studies Hybridoma Bank (Iowa City, IA). anti-RFP (5F8) from ChromoTek (Munich, Germany).

### Colony formation

Colony formation was performed as described previously(25). On each well of six-well plates, 1,000 cells were seeded and grown in 2 ml DMEM media for 14 days or 10,000 cells were seeded and grown for ten days. The colonies were fixed with 4% formaldehyde and then stained with 0.5% crystal violet. Images were taken using ChemiDoc Touch Imager (Bio-Rad, Hercules, CA) and area of colonies were determined by image J software (National Institutes of Health). Three biological replicates were analyzed.

### Western blotting

Cells were washed with PBS, and cell lysates were prepared in cold Triton lysis buffer (1% Triton X-100; 40 mm Hepes, pH 7.5; 120 mm sodium chloride; 1 mm EDTA; 1 mm phenylmethylsulfonyl fluoride; 10 mm sodium pyrophosphate; 1 μ;g/ml each of chymostatin, leupeptin, and pepstatin; 10 μ;g/ml each of aprotinin and benzamidine; 2 μg/ml antipain; 1 mm sodium orthovanadate; and 50 mm sodium fluoride). cell lysates were centrifuged to remove cell debris. Protein concentration was determined using the Bio-Rad Protein Assay (Bio-Rad) with BSA as standard then were prepared in Laemmli SDS-PAGE sample buffer. Aliquots of protein lysate were resolved by SDS-PAGE, transferred to nitrocellulose membranes (0.45 μm) (Bio-Rad) and immunoblotted with the indicated antibodies, followed by horseradish peroxidase-conjugated goat anti-mouse or goat anti-rabbit IgG (Sigma-Aldrich) and Amersham ECL Select Western Blotting Detection Reagent or Pierce ECL Western Blotting Substrate (Thermo Scientific, Hudson, NH). Signals were detected using ChemiDoc Touch Imager. For Western blot signal quantitation, the Image Lab software (Bio-Rad) was used.

### Biochemical subcellular fractionation

Subcellular fractionation was performed as described previously(6). After rinsing with cold PBS, cold lysis buffer (20 mM HEPES pH 8.0, 1 mM EDTA, 1.5 mM MgCl_2_, 10 mM KCl, 1 mM DTT, 1 mM sodium orthovanadate, 1 mM NaF, 1 mM PMSF, 0.5 mg/mL benzamidine, 0.1 mg/ml leupeptin, and 1.2 mg/mL aprotinin) was used to lyse the cells. Then 7.5 μL of 10% NP-40 detergent was added to the cells after incubating them on ice for 15 min. right after adding NP-40 detergent to the cells. Lysates were centrifuged for 1 min at 14000 g and 4°C, and supernatants were collected as cytosolic fractions. Pellets were washed 4 times and incubated for 40 min at 4°C (1% Triton X-100, 40 mM HEPES (pH 7.5), 120 mM sodium chloride, 1 mM EDTA, 1 mM phenyl methylsulfonyl fluoride, 10 mM sodium pyrophosphate, 1 μg/ml each of cymostatin, leupeptin and pepstatin, 10 μg/ml each of aprotinin and benzamidine, 2 μg/ml antipain, 1 mM sodium orthovanadate, 50 mM sodium fluoride). Supernatants were collected as nuclear fractions after centrifugation for 10 min at 13,800 g and 4°C.

### Co-immunoprecipitation

Cells were washed with PBS and cell lysates prepared in cold triton lysis buffer (1% Triton X-100; 40 mm HEPES (pH 7.5); 120 mm sodium chloride; 1 mm ethylene diamine-tetraacetic acid; 1 mm phenyl methylsulfonyl fluoride; 10 mm sodium pyrophosphate; 1 μg/ml each of cymostatin, leupeptin, and pepstatin; 10 μg/ml each of aprotinin and benzamidine; 2 μg/ml antipain; 1 mm sodium orthovanadate; 50 mm sodium fluoride) supplemented with 2% empigen for anti-K19 IP, or cold NP-40 lysis buffer (0.25% NP-40; 50 mM Tris (pH 8.0); 100 mM sodium chloride; 1 mm phenyl methylsulfonyl fluoride; 10 mm sodium pyrophosphate; 1 μg/ml each of cymostatin, leupeptin, and pepstatin; 10 μg/ml each of aprotinin and benzamidine; 2 μg/ml antipain; 1 mm sodium orthovanadate; 50 mm sodium fluoride) for anti-HNRNPK IP. Cell lysates were centrifuged to remove cell debris, and protein concentration was determined using the Bio-Rad Protein Assay with BSA as standard. Aliquots of cell lysate were then incubated with the indicated antibody or IgG control, and immune complexes were captured using Protein G Sepharose (GE Healthcare).

### Immunofluorescence (IF) staining

IF staining of cells was performed as described previously(26). Cells grown on glass coverslips (VWR, Radnor, PA) were washed with 1X PBS, fixed in 4% paraformaldehyde in 1X PBS for 35 min, and permeabilized in 0.1% Triton X-100 for 20 min or 0.01 Digitonin for 5 minutes. Samples were blocked in 5% normal goat serum (NGS; RMBIO, Missoula, MT) in 1X PBS before staining with primary antibodies diluted at 1:400 ratio in 5% NGS blocking buffer and a mixture of 1:1000 of Alexa Fluor 488-conjugated goat anti-mouse secondary antibody (Invitrogen) and 1:5000 DAPI (Sigma-Aldrich in 1X PBS) was added for 1 h incubation at RT. After 1X PBS washes, coverslips were mounted on microscope slides with mounting medium containing 1,4-diaza-bicyclo[2.2.2]octane (Electron Microscopy Sciences, Hatfield, PA). Fluorescence images were taken using the Olympus optical elements fluorescence microscope (Olympus Optical Co., Japan).

### Proximity ligation assay

The Duolink in situ proximity ligation assay (PLA) was performed according to the manufacturer’s protocol (Sigma–Aldrich). In brief, cells were plated on glass coverslips, rinsed three times with PBS and fixed in 3.7% formaldehyde in PBS for 20 min. The cells were permeabilized in 0.01% digitonin for 5 min and blocked with 5% NGS in PBS for overnight at 4 °C. After blocking, cells were then incubated with antibodies against HNRNPK, K19 and IgG in PBS containing 5% NGS overnight at 4 °C, followed by incubation with corresponding secondary antibodies conjugated with PLA probes for 60 min at 37 °C in the dark. Cells were washed three times in PBS. Duolink and DAPI signals were detected using Olympus optical elements fluorescence microscope (Olympus). Images are analyzed using ImageJ.

### Graphs and statistics

All graphs in the manuscript are shown as mean ± standard error of mean. For comparisons between two datasets, a Student’s t test (tails = 2, type = 1) was used, and statistically significant p-values are indicated in the figures and figure legends.

## Results

### K19 knockout reduces proliferation of MDA-MB-231 cells

We set out to study the effect of K19 on HNRNPK localization and posttranscriptional gene regulatory activity in a cell culture model system. We used MDA-MB-231 triple negative breast cancer (TNBC) cells with CRISPR-Cas9 mediated *KRT19* knockout (KO)(26) and confirmed complete ablation of K19 expression by Western blotting (Supplementary Figure 1a) and RNA-sequencing (RNA-seq) (Supplementary Table 1).

A previous study reported that loss of K19 in estrogen receptor-positive MCF7 breast cancer cells resulted in a delayed cell cycle(29). We tested whether K19 also affected proliferation of TNBC cells by cell counting and found that *KRT19* KO cells indeed showed significantly decreased cell proliferation compared to the parental control (Figure 1a). Similarly, *KRT19* KO cells showed significantly reduced cell proliferation in a colony formation assay (Figure 1b). We confirmed the specificity of the *KRT19* KO effect on cell proliferation by rescuing K19 expression with a GFP-K19 construct (Supplementary Figure 1b,c) in our cells, which as expected, resulted in increased cell proliferation by cell counting (Figure 1d) and colony formation assays (Figure 1e, f). Overall, our findings show that K19 promotes TNBC cell proliferation.

**Figure 1.**
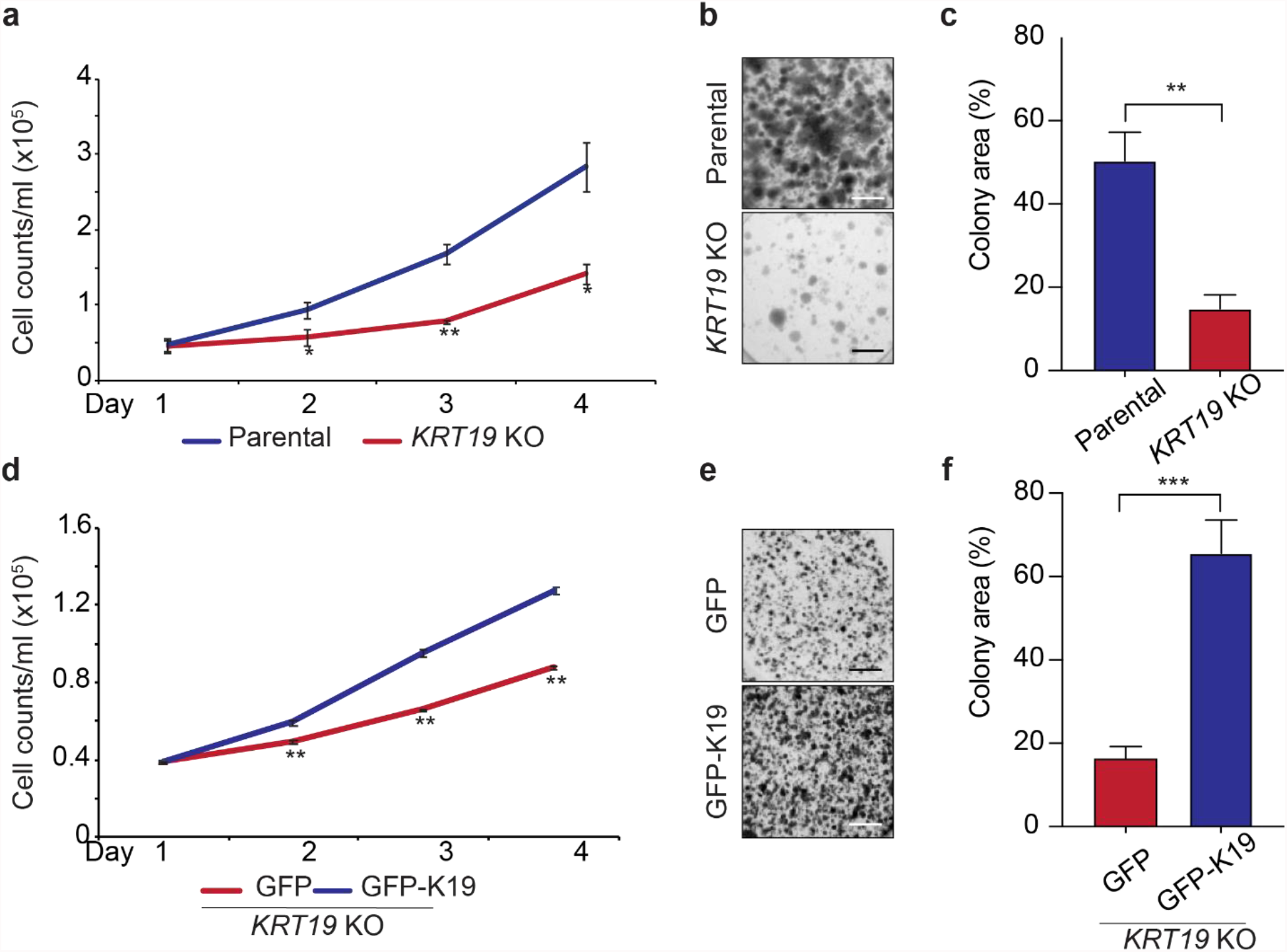
Cell proliferation of MDA-MB-231 breast cancer cells requires K19. **a)** Cells were counted every 24 h for 4 days after initial seeding of 50,000 parental (blue line) or *KRT19* KO cells (red line). **b)** Colony formation assay. Two weeks after seeding 1000 cells in a 6 well plate-format, cells were stained by crystal violet to visualize colonies, and plates were imaged and **c)** colony area quantified using the ImageJ software. **d)** Same as in a) using except using *KRT19* KO cells stably expressing either GFP (red line) or GFP-K19 (blue line). **e, f)** Same as in b) and c) except using *KRT19* KO cells stably expressing GFP or GFP-K19. Data from at least three biological replicates are shown as mean ± standard error of mean. * p < 0.05, ** p < 0.01, *** p < 0.001. Scale bar 0.5 mm.

### K19 binds to HNRNPK and promotes its cytoplasmic localization

Keratins are an interaction platform for various proteins(30), and a previous study showed that a keratin closely related to K19, K17, interacts with HNRNPK in skin cancer cells(6). To test a potential interaction between K19 and HNRNPK, we performed proximity ligation assay (PLA). PLA using K19 and HNRNPK antibodies showed fluorescence signals from parental cells but not from *KRT19* KO cells, suggesting that K19 interacts with HNRNPK (Figure 2a). Note that we focused on the cytoplasm in our PLA by permeabilizing with mild detergents (0.01% digitonin) to avoid saturation of available antibody by the predominantly nuclear fraction of HNRNPK. We validated the K19-HNRNPK interaction by co-immunoprecipitation (co-IP) and found that K19 and HNRNPK were detected in IPs of HNRNPK and K19, respectively (Figure 2b). Consistently, in HEK293 cells HNRNPK specifically co-immunoprecipitated with transiently expressed GFP-tagged K19 transiently expressed but not in the controls (Figure 2c).

**Figure 2.**
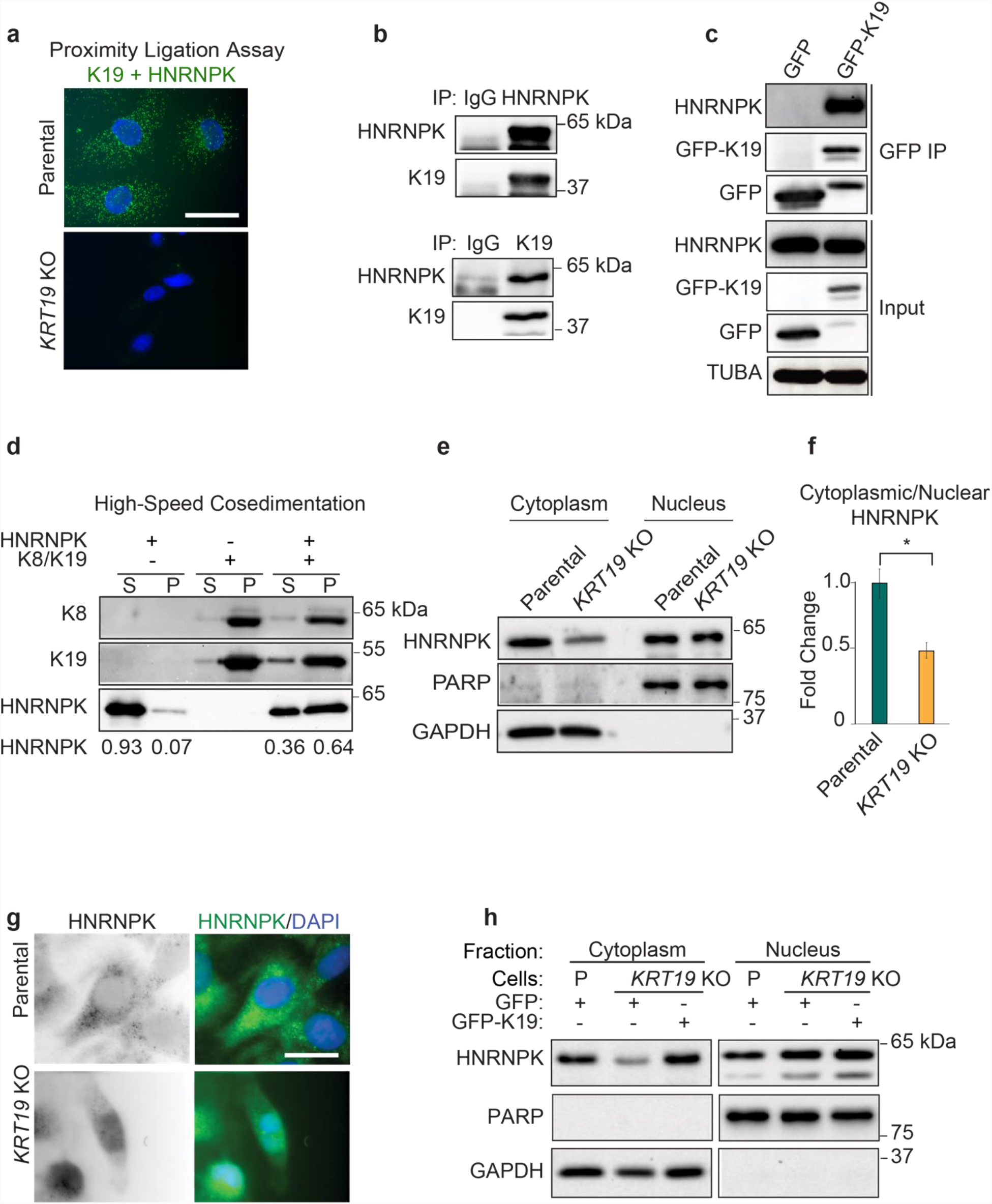
K19 binds to hnRNPK and promotes the cytoplasmic localization of HNRNPK. Proximity ligation assay using anti-K19 and HNRNPK antibodies in MDA-MB-231 parental and *KRT19* KO cells. Bar = 10 μm. **b)** Co-IP of endogenous K19 and HNRNPK in MDA-MB-231 cells using IgG as a control. **c)** IP’s using anti-GFP antibody in HEK293 cells transfected with GFP or GFP-K19 were immunoblotted with antibodies against the indicated proteins. **d)** High-speed co-sedimentation assay following recombinant HNRNPK incubation with (+) or without (-) preassembled K8/K19 filaments. Supernatant (S) and pellet (P) were separated, and immunoblotting performed. **e)** Biochemical subcellular fractionation of parental and *KRT19* KO cells. Cytoplasmic and nuclear fractions were immunoblotted with antibodies against the indicated proteins and **f)** quantified (data from at least three biological replicates are shown as mean ± standard error of mean). **g)** Parental and *KRT19* KO cells permeabilized using 0.01% Digitonin were immunostained with anti-HNRNPK antibody. Nuclei were stained with DAPI (bar = 10 μm). **h)** Biochemical subcellular fractionation of parental or *KRT19* KO cells stably expressing GFP or GFP-K19. Cytoplasmic and nuclear fractions were immunoblotted with antibodies against the indicated proteins.

We then tested direct interaction between K19 filaments and HNRNPK *in vitro* by high-speed co-sedimentation. For this, filament was preassembled using purified K19 and its oligomerization partner K8. Next, recombinant HNRNPK was incubated with or without preassembled K8/K19 filaments. After high-speed centrifugation, sedimentation of recombinant HNRNPK increased significantly in the presence of K8/K19 filaments, whereas HNRNPK remained mostly in the supernatant fraction without filaments (Figure 2d), indicating a direct interaction between HNRNPK and K19 filaments.

Keratin filaments regulate the subcellar localization of several proteins(6), and we hypothesized that K19 may also influence localization of the typically predominantly nuclear HNRNPK. Therefore, we examined the cytoplasmic and nuclear levels of HNRNPK in parental and *KRT19* KO cells. Biochemical subcellular fractionation showed that there is a decrease in cytoplasmic HNRNPK level and cytoplasmic to nuclear ratio of HNRNPK in *KRT19* KO cells (Figure 2e, f) without changes in total levels of HNRNPK (Supplementary Figure 2b). Immunostaining of cytoplasmic HNRNPK after mild digitonin permeabilization also showed higher HNRNPK signal from the cytoplasm of parental MDA-MB-231 cells compared to *KRT19* KO cells (Figure 2h). Finally, we confirmed the requirement of K19 on cytoplasmic HNRNPK accumulation by assessing its level in *KRT19* KO cells after stable expression of a GFP-K19 rescue construct (Figure 2g). In summary, we concluded that K19 directly interacts with and regulates cytoplasmic localization of HNRNPK.

### K19 is required for the stability of cytoplasmic HNRNPK targets

As an RNA binding protein (RBP), HRNPK is expected to posttranscriptionally regulate its RNA targets in the cytoplasm(15) and therefore we examined whether loss of K19 and concomitant loss of cytoplasmic HNRNPK resulted in changes in RNA regulation. First, we tested the effect of K19 on the HNRNPK mRNA target repertoire. We comprehensively mapped the RNA interactome of HNRNPK and characterized its RNA recognition elements (RREs) using 4-thiouridine (4SU) PAR-CLIP in both parental and *KRT19* KO cells(27). In these cells, RNA and proteins were crosslinked by UV, HNRNPK immunoprecipitated, and the RNA covalently attached in immunoprecipitated ribonucleoproteins (RNPs) was partially digested and radiolabeled. Autoradiography of the HNRNPK immunoprecipitate revealed a single band migrating at ∼60 kDa in both cell lines, corresponding to the HNRNPK-RNP (Figure 3a). We recovered and sequenced HNRNPK-bound RNA fragments from parental MDA-MB-231 cells and *KRT19* KO cells and used PARalyzer to identify clusters of overlapping reads harboring the characteristic T-to-C mutations indicating RNA-protein crosslinking events(31). Overall, we identified 75,557 and 24,874 HNRNPK binding sites on the whole genome in parental and *KRT19* KO cells, respectively (Supplementary Tables 2 and 3). RNAs binding to HNRNPK more in parental cells included transcripts known to bind to HNRNPK such as those coding for EGFR or HNRNPK itself (6). Using MEME, we found that our PAR-CLIP binding sites were highly enriched for C/U-rich sequences, consistent with the previously reported HNRNPK RRE(12) (Figure 3b, c). As expected from a predominantly nuclear protein, most HNRNPK binding sites localized to pre-mRNA introns, nevertheless, we found a prominent contribution of the 3’ untranslated regions (UTR) to the binding profile (Figure 3d), which could result from the contribution of cytoplasmic HNRNPK.

**Figure 3.**
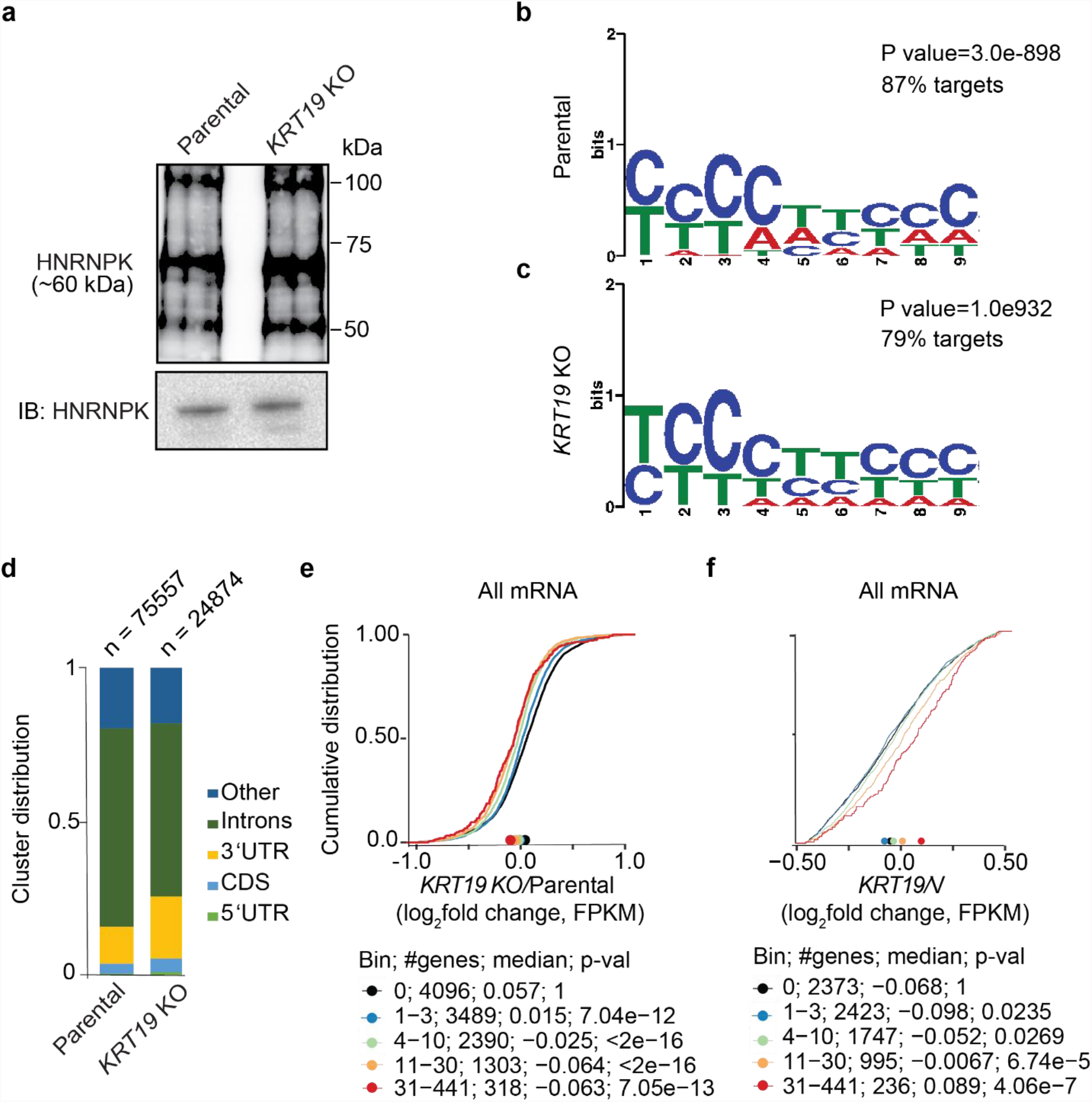
K19 stabilizes mRNAs directly bound to HNRNPK. **a)** Autoradiograph of crosslinked and radiolabelled HNRNPK separated by SDS-PAGE and immunoblot of HNRNPK from parental and *KRT19* KO cells used for PAR-CLIP. **b**,**c)** Sequence logo representation of the HNRNPK in b) parental and c) *KRT19* KO cells. **d)** Distribution of crosslinked sequence reads from PAR-CLIP of HNRNPK in parental and *KRT19* KO cells. **e**,**f)** mRNA expression changes upon *KRT19* KO in MDA-MB-231 cells determined by RNAseq (n=3 biological triplicates). The empirical cumulative distribution function (CDF) of HNRNPK targets in **e)** *KRT19* KO/parental cells and **f)** cells stably overexpressing *KRT19*/vector (V) binned by number of binding sites compared to expressed non-targets. Targets (colored lines) binned by number of HNRNPK binding sites and non-targets (black line) with minimal gene expression of 4 fragments per kilobase of exon per million mapped fragments ((FPKM) ≥4) are shown.

We argued that the loss of cytoplasmic HNRNPK in *KRT19* KO cells may result in changes in posttranscriptional gene regulation, in particular mRNA stability and abundance. Thus, we next examined the role of K19 on the stability of HNRNPK targets and integrated RNA-seq results from parental and *KRT19* KO cells (Supplementary Table 1) with our PAR-CLIP data. We found that *KRT19* KO and therefore reduction of cytoplasmic HNRNPK results in a reduction of HNRNPK target mRNA abundance, with the magnitude of this effect depending on the overall strength of HNRNPK binding as defined by the number of HNRNPK binding sites (Figure 3e). In particular mRNAs with HNRNPK-binding sites at their 3’ UTR - a region that we suspect is bound by cytoplasmic HNRNPK - were destabilized in *KRT19* KO cells (Supplementary Figure 3a). mRNAs with HNRNPK-binding sites at 5’UTRs and coding regions were not destabilized in *KRT19* KO cells (Supplementary Figure 3c, d). Interestingly, mRNAs with HNRNPK-binding sites in introns were also dependent on K19 (Supplementary Figure 3b), though these same mRNAs also contained HNRNPK 3’UTR binding sites, opening up the possibility that HNRNPK generally binds all its targets first in the nucleus and then shuttles as part of a larger RNP into the cytoplasm.

Increase of K19 expression levels by stable overexpression (Supplementary Figure 3e) further increased abundance of HNRNPK targets, likely by increasing cytoplasmic HNRNPK levels (Figure 3f). Again, this effect was most pronounced in the strongest HNRNPK targets, as well as those bound by HNRNPK in 3’UTR, but was not detectable when HNRNPK targets were binned by binding in coding regions or 5’UTR (Supplementary Figure 3f-i). In summary, our data suggest that K19-dependent cytoplasmic localization of HNRNPK results in increased target mRNA abundance, possibly by competition with other, destabilizing RBPs.

### Re-expression of cytoplasmic HNRNPK in KRT19 KO cells rescues decreased HNRNPK target stability

We then examined whether overexpressing cytoplasmic HNRNPK in *KRT19* KO cells can rescue the destabilization of HNRNPK targets due to the absence of K19. To increase cytoplasmic levels of HNRNPK we created an expression construct where nuclear localization signal was deleted (HNRNPK ΔNLS) which localized to the cytoplasm (Figure 4a-c). Integrating the list of HNRNPK targets (Supplementary Table 2) with RNA-seq results (Supplementary Table 4) from *KRT19* KO cells overexpressing HNRNPK ΔNLS or vector showed that HNRNPK ΔNLS overexpression revealed an increase in abundance of HNRNPK target mRNAs, independent of whether the targets were binned by overall number of HNRNPK binding sites or by number of binding sites in the target mRNA 3’UTR (Figure 4d, e). This indicated that the negative effect of K19 loss on HNRNPK targets was indeed due to loss of cytoplasmic HNRNPK.

**Figure 4.**
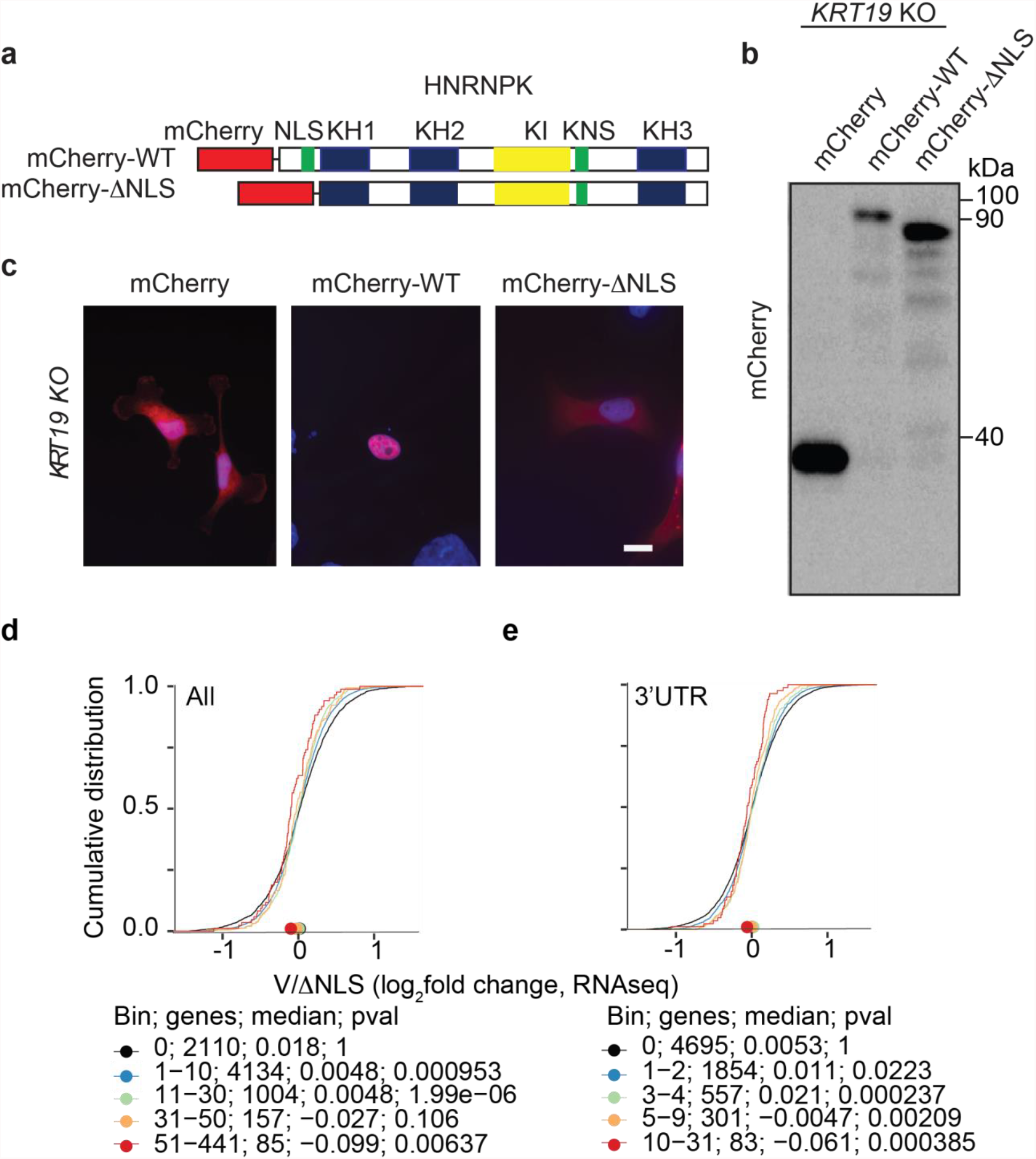
Overexpressing cytoplasmic HNRNPK rescued the stability of targets in *KRT19* KO cells. **a)** Schematics of wildtype (WT) HNRNPK and cytoplasmic mutant (ΔNLS). N-termini of WT and ΔNLS HNRNPK were tagged with mCherry (mCherry-WT and mCherry-ΔNLS, respectively). **b)** Immunoblot from lysates of *KRT19* KO cells transfected with either mCherry, mCherry-WT, or mCherry-ΔNLS using an antibody against mCherry. **c)** *KRT19* KO cells transiently transfected with mCherry, mCherry-WT, or mCherry-ΔNLS were immunostained with anti-RFP antibody. Nuclei are shown with DAPI (bar = 20 μm). **d**,**e)** The empirical cumulative distribution function of mRNA expression changes in *KRT19* KO cells transiently transfected with mCherry or mCherry-ΔNLS with genes binned by HNRNPK binding **d)** across entire transcript or **e)** in the 3’ UTR. Targets (colored lines) binned by number of HNRNPK binding sites and non-targets (black line) with minimal gene expression of 4 fragments per kilobase of exon per million mapped fragments ((FPKM) ≥4) are shown.

### K19 and HNRNPK co-regulate p53 signaling pathway

We next identified molecular pathways regulated by HNRNPK that are dependent on K19 to find gene networks that may give insight into the apparent association of K19 loss with cell proliferation. First, we analyzed gene ontology (GO) pathways that significantly changed upon HNRNPK KD (Supplementary Table 5), focusing on downregulated transcripts (log_2_ fold change > 0.4), considering that we found that HNRNPK stabilizes its targets. The p53 signaling pathway was the most significantly affected in HNRNPK KD cells (Figure 5a). We further narrowed the list of input genes into GO analysis by only considering top HNRNPK targets that are downregulated (≥ 6 PAR-CLIP binding sites, log_2_ fold change > 0.4, Supplementary Table 6) and again found that p53 signaling was among the three most enriched terms (Figure 5b). Finally, we repeated this analysis using transcriptome data from MDA-MB-231 cells after *KRT19* KO, which we showed previously selectively affects HNRNPK targets bound in the cytoplasm. Again, and consistent with the HNRNPK KD data, p53 signaling emerged among the top three pathways with downregulated HNRNPK targets (Figure 5c, d) in addition to pathways related to cell growth and proliferation, as previously observed in MCF-7 cells after *KRT19* KO(29).

**Figure 5.**
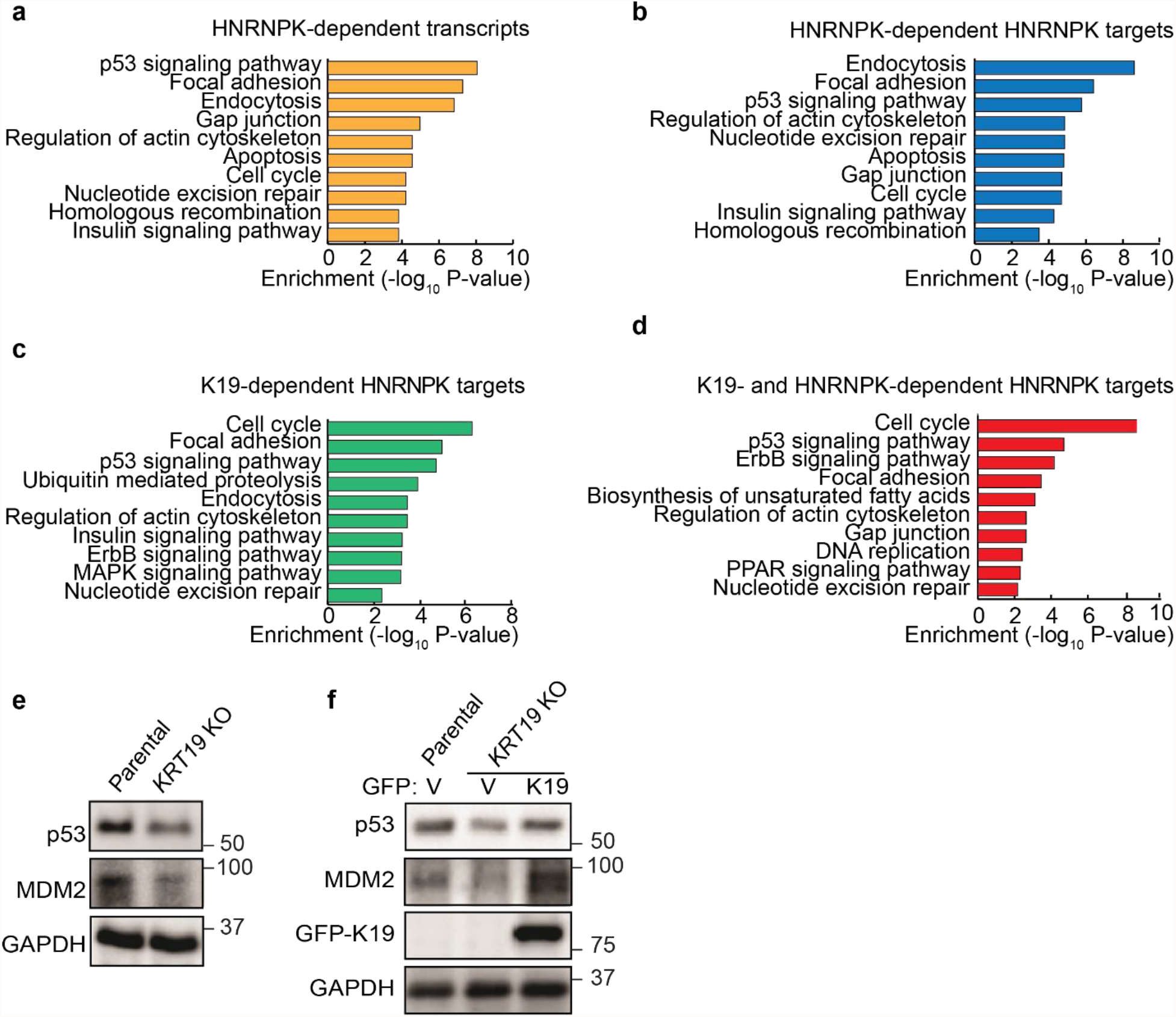
Gene ontology identifies that K19 and HNRNPK promotes p53 signaling pathway in MDA-MB-231 cells. **a)** Enrichment of Gene Ontology (GO) terms among mRNAs downregulated by HNRNPK KD. **b**,**c)** Enriched GO terms among downregulated HNRNPK PAR-CLIP targets after b) HNRNPK KD or c) K19 KO. **d)** GO terms enriched among HNRNPK target mRNAs downregulated in both K19 KO and HNRNPK KD data. GO terms for top 10 pathways based on p values are shown. **e**,**f)** Immunoblots from lysates of e) parental MDA-MB-231 and *KRT19* KO cells or f) parental and *KRT19* KO cells stably expressing GFP or GFP-K19 with antibodies against the indicated proteins.

Dysregulation of the p53 pathway by loss of expression or mutation of its components is one of the major drivers promoting cancer cell survival and metastasis and our TNBC MDA-MB-231 cell line expresses high levels of the p53 R280K mutant(32). We decided to test whether K19 loss changes expression levels of p53 itself or its activator MDM2. We found that p53 and MDM2 levels are decreased in *KRT19* KO cells (Figure 5e). This decrease was dependent on K19, as reintroduction of GFP-K19 rescued p53 and MDM2 expression levels (Figure 5f). Altered p53-pathway gene expression may thus contribute to the observed decreased cancer cell proliferation accompanying K19 loss.

## Discussion

Keratin 19 plays a role in posttranscriptional gene regulation by affecting subcellular localization of HNRNPK, thereby altering expression of HNRNPK target mRNAs bound to HNRNPK via 3’UTR (Figure 6). We showed that HNRNPK and K19 physically interact in the cytoplasm, and K19 and HNRNPK co-regulate p53 signaling pathway. Our findings identify a comprehensive list of HNRNPK-dependent mRNAs and reveal a novel gene regulatory mechanism for HNRNPK.

**Figure 6.**
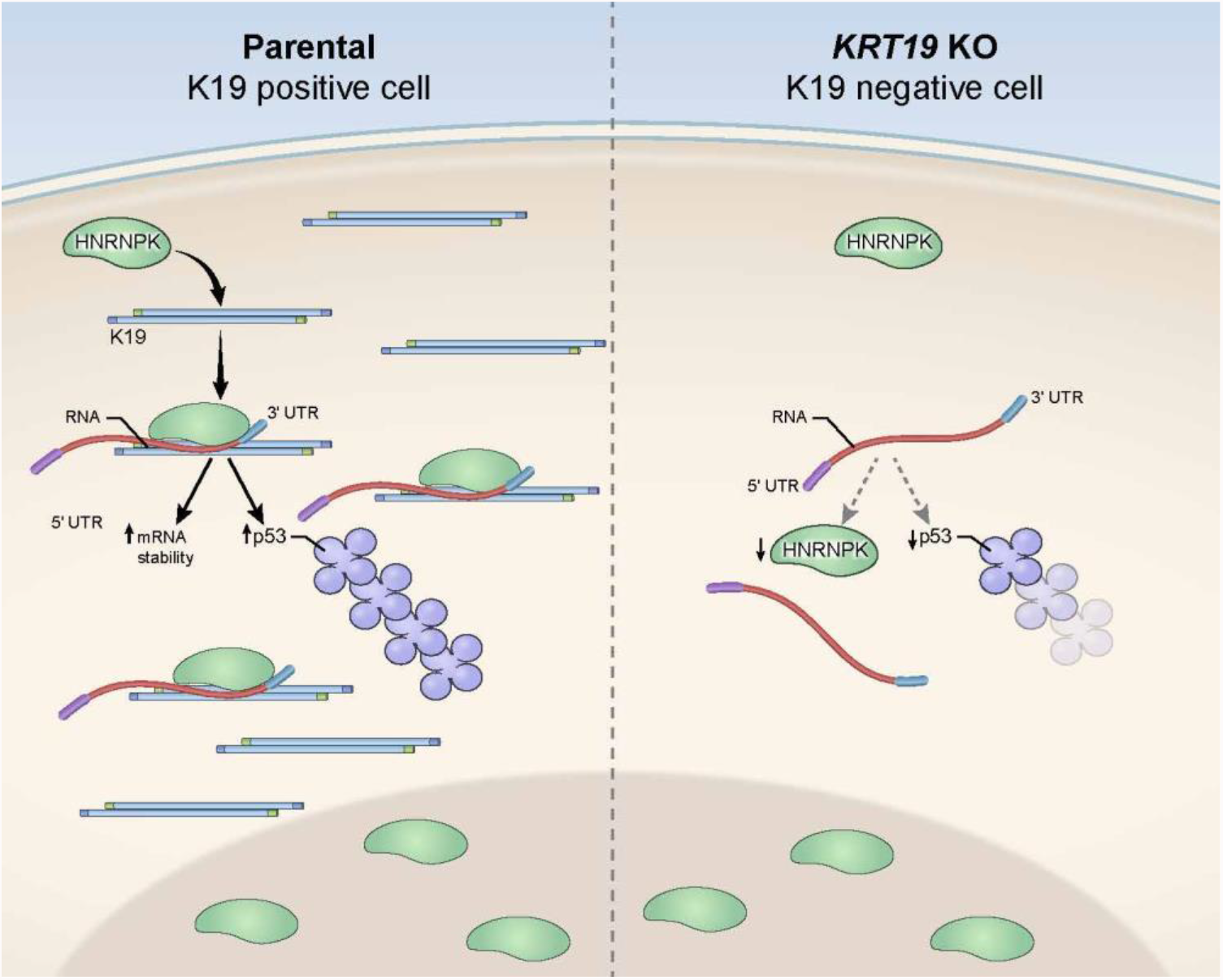
A model of how K19 regulates HNRNPK. Left panel: K19 binding to HNRNPK retains HNRNPK in the cytoplasm. In the cytoplasm, HNRNPK binds to 3’ UTR of its target mRNAs to stabilize them. Targets stabilized by HNRNPK regulate various signaling pathways including p53 signaling pathway. Right panel: in the absence of K19, decreased level of HNRNPK in the cytoplasm destabilizes HNRNPK-bound mRNAs.

In both parental and *KRT19* KO cells, HNRNPK binding preferences were not altered and it recognized the expected C-rich sequences(12,33). This suggested that K19 has little to no effect on the mechanism of nucleic acid recognition by HNRNPK and that changes in HNRNPK gene regulatory activity were most likely due to changes in its localization. Indeed, keratins regulate the localization of proteins in the cytoplasm to impact cellular and physiological processes including cell cycle, translation, cell-cell adhesion(25,34–36). Our data show that in the presence of K19, HNRNPK increases the abundance of target mRNAs in the cytoplasm, very likely by increasing their stability and possibly through competition with other RBPs(37). Testing half-lives of 3’ UTR targets of HNRNPK will be required to confirm an influence of K19 on stability of HNRNPK targets. Currently, the exact mechanism of how K19 retains HNRNPK in the cytoplasm remains unclear. Since HNRNPK is a nucleocytoplasmic shuttling protein(17), interaction between K19 and HNRNPK in the cytoplasm and HNRNPK co-sedimentation with K8/K19 filaments indicate that HNRNPK becomes associated with K8/K19 filaments in the cytoplasm after exiting the nucleus. Thus, K19-containing filaments would serve as cytoplasmic loading docks for HNRNPK. Due to its role in RNA transport(38,39), it is likely that HNRNPK emerges from the nucleus already bound to its targets. Under this scenario, the interaction with K19 would allow HNRNPK to better retain and stabilize targets. Preloading of HNRNPK to its preferred targets may also explain why overall number of PAR-CLIP binding sites was predictive of HNRNPK-mediated regulation of target RNA stability in the cytoplasm, even if they were not exclusively found in the cytoplasm, but additionally in introns.

It will be important to explore whether HNRNPK shows better affinity to filamentous or soluble K19. In this context, posttranslational modifications play a major role in keratin filament dynamic. Compared to K19 proteins in filaments, soluble K19 proteins are hyperphosphorylated and SUMOylated(40). Indeed, the K17-HNRNPK interaction in keratinocytes required intact Ser44 of K17, a RSK-dependent phosphorylation site(6). This serine residue is conserved across type I epithelial keratins, and K19 has its own at amino acid 35 position(41,42). Ser also happens to be a major phosphorylation site of K19 and has been shown to be phosphorylated by Akt(42,43). Other signaling pathways such as TGF-β(44) and Erk1/2(18) which affect the cytoplasmic accumulation of HNRNPK may also be involved in regulating RNA stabilizing function of HNRNPK by K19. While this is not our favored interpretation, there exists a possibility that K19 interacts with HNRNPK inside the nucleus. Keratins have been found inside the nucleus, where they interact with nuclear proteins(45). In addition, K19 contains a bipartite nuclear localization signal towards its C-terminus(45). Regardless of the exact location of their interaction however, it is likely that K19-bound HNRNPK represents only a small minority of the overall pool. Most of HNRNPK resides inside the nucleus while most of K19 is in the cytoplasm. Nevertheless, destabilization of HNRNPK targets upon K19 demonstrates the profound effect K19 has on HNRNPK and likely the gene regulatory importance of cytoplasmic HNRNPK(6).

Top 10 pathways involving HNRNPK targets identified to be dependent on K19 involved regulating cancer progression such as cell cycle, p53 and ErbB signaling. Our observation of p53 signaling pathway is consistent with previous reports that HNRNPK serves as a p53 partner and regulator in response to stress(11,46). Of note, MDA-MB-231 cells, like 80% of TNBCs, express an oncogenic mutant p53 which promotes cell survival and tumorigenesis(47,48). Also, although MDM2 is a ubiquitin ligase which targets p53 for degradation, MDM2 in TNBC can promote metastasis and its increased level is associated with poor patient prognosis(49). Identification of genes involved in promoting cancer is a strong indication that transcriptomic changes of those genes significantly contribute towards decreased cell proliferation of *KRT19* KO cells. Therefore, we have not only identified novel RNA targets of HNRNPK in breast cancer cells, but also how they are stabilized.

The fact that K19 was required for the cytoplasmic localization of HNRNPK further suggests that K19 and HNRNPK interaction contributes towards cancer progression and metastasis. Indeed, higher levels of K19 as well as cytoplasmic localization of HNRNPK in tumors has been correlated with poor patient survival rate and metastasis(7). Therefore, our data demonstrates a potential explanation of these clinical observations. Given the fact that both K19 and HNRNPK are members of large families of proteins, it is likely that there are other intermediate filament protein-HNRNP interactions. Indeed, K17 interacts with HNRNPK as mentioned above, and HNRNPS1 has been shown to interact with a type III intermediate filament protein vimentin, which is upregulated in several cancers and plays a critical role in metastasis(50). Therefore, our findings can be used to characterize the mechanism underlying progression of various tumor types.

## Acknowledgements

We thank past and current members of the Coulombe, Chung, and Hafner labs for their support and helpful comments. This work was supported by National Cancer Institute grant R15CA2113071 (to BMC) and National Institute for Arthritis, Musculoskeletal and Skin diseases grant R01AR044232 (to PAC). We thank the NIAMS Genomics, Light Imaging Section, and Flow Cytometry Core Facilities and Gustavo Gutierrez-Cruz, James Simone, Jiff Lay, Kevin Tinsley, Faiza Naz, and Drs. Stefania Dell’Orso and Davide Randazzo (NIAMS/NIH) for sequencing and flow cytometry support.

## Author contributions

A.F., B.M.C., and M.H. conceived the experiments. A.F., P.S., X.W, A.E. conducted the experiments. A.F., D.G.A., A.M., A.J., S.A., M.H., and B.M.C. analyzed the results. B.M.C., P.A.C. and M.H. provided resources and guidance. A.F., B.M.C., and M.H. wrote the manuscript. All authors reviewed the manuscript.

## Notes

**Competing interests:** The authors declare no conflict of interest.

### Competing Interest Statement

The authors have declared no competing interest.

