## Supplemental Information for "HNRNPK is retained in the cytoplasm by Keratin 19 to stabilize target mRNAs"

### Supplementary Figures

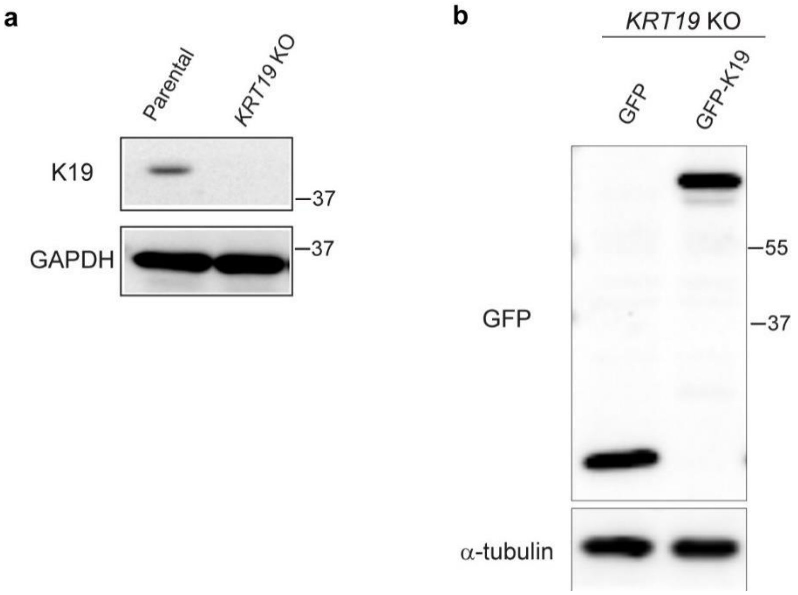


**Supplementary Figure 1.** K19 levels in KO and rescue cell lines. **a)** Whole cell lysates of parental control and *KRT19* KO cell lines were harvested, and immunoblotting was performed with antibodies against the indicated proteins. **b)** Whole cell lysates of *KRT19* KO cell lines stably expressing either GFP or GFP-K19 were harvested, and immunoblotting was performed with antibodies against the indicated proteins.

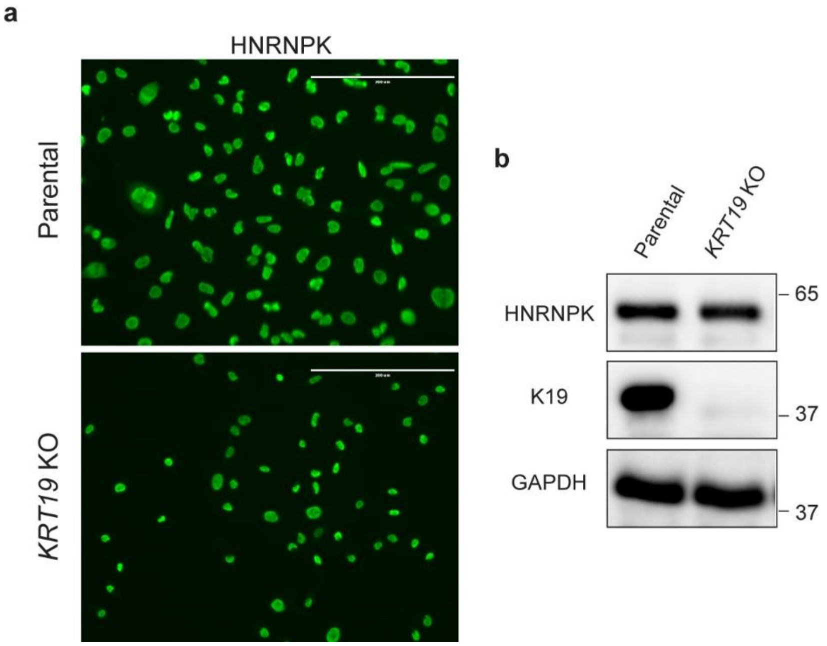


**Supplementary Figure 2.** Total HNRNPK levels in parental and *KRT19* KO MDA-MB-231 cells do not differ. **a)** Parental and *KRT19* KO cells immunostained with anti-HNRNPK antibody (bar = 200 µm). **b)** Immunobloy for indicated proteins from whole cell lysates of parental and *KRT19* KO cells.

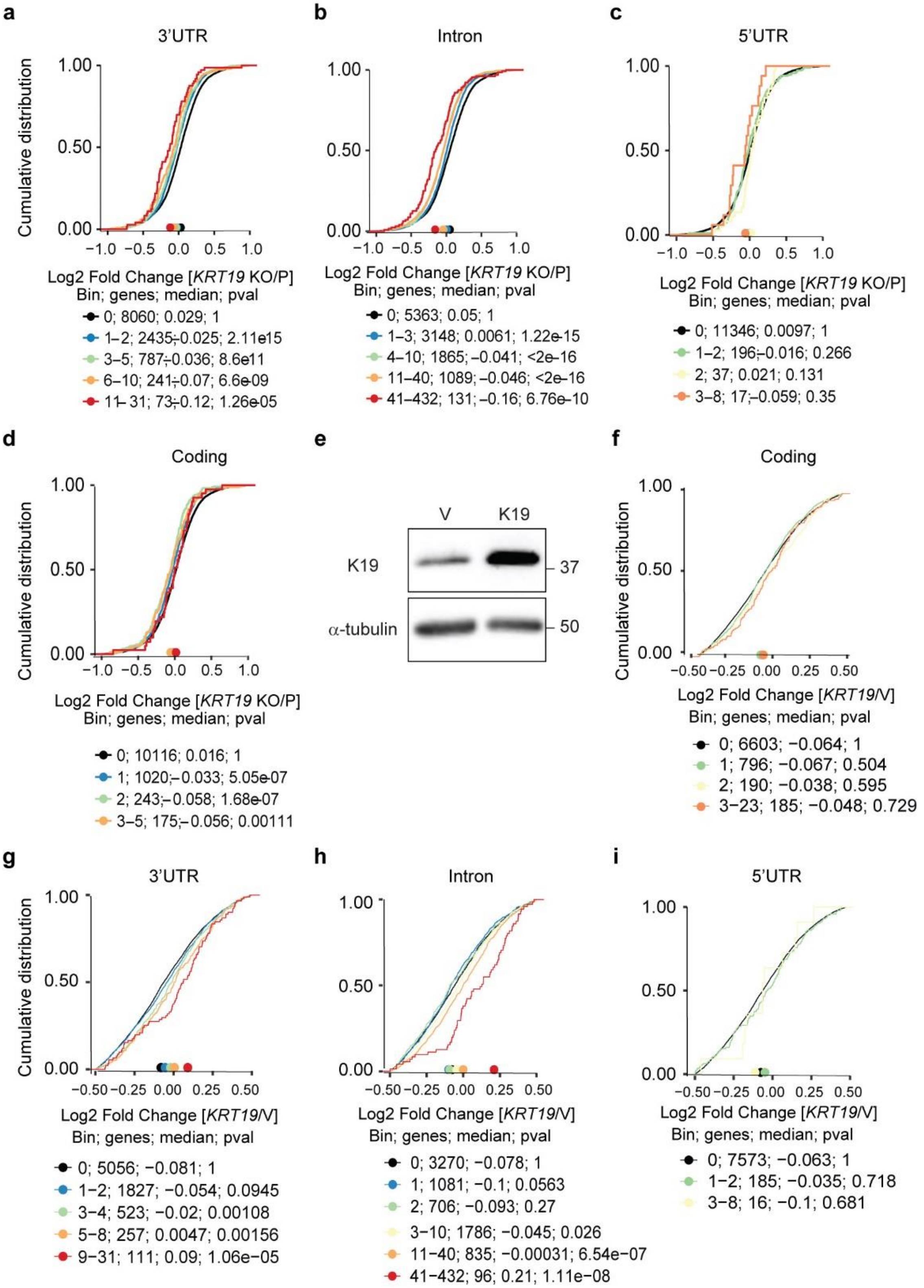


**Supplementary Figure 3. K19 specifically destablized mRNAs bound to HNRNPK via introns and 3' UTR.** The empirical cumulative distribution function of mRNA expression changes upon *KRT19* KO. mRNAs bound to HNRNPK were binned by number of **a)** 3’ UTR, **b)** intron, **c)** 5’UTR and **d)** coding region in *KRT19* KO and parental cells. e. Immunoblot from lysates of MDA-MB-231 cells stably expressing vector or K19 were using antibodies against the indicated proteins. The empirical cumulative distribution function of mRNA expression changes in MDA-MB-231 cells overexpressing vector or *KRT19.* mRNAs bound to HNRNPK were binned by number of binding sites in the **f)** coding region, **g)** 3’ UTR, **h)** intron, and **i)** 5’UTR in. Only mRNAs with expression of FPKM ≥4 are shown. Targets (colored lines) binned by number of HNRNPK binding sites and non-targets (black line) with minimal gene expression of 4 fragments per kilobase of exon per million mapped fragments ((FPKM) ≥4) are shown.
